# Background matching can reduce responsiveness of jumping spiders to stimuli in motion

**DOI:** 10.1101/2023.05.09.539969

**Authors:** Min Tan, Jeremiah Y.O. Chan, Long Yu, Eunice J. Tan, Daiqin Li

**Affiliations:** Department of Biological Sciences, National University of Singapore, 14 Science Drive 4, Singapore 117543; Centre for Behavioural Ecology & Evolution, College of Life Sciences, Hubei University, Wuhan 430062, Hubei, China; Division of Science, Yale-NUS College, 16 College Avenue West, Singapore 138527

**Keywords:** Background matching, Camouflage, Motion, Jumping spiders, Salticidae, Visual acuity, Visual responsiveness

## Abstract

Motion and camouflage were previously considered to be mutually exclusive, as sudden movements can be easily detected. Background matching, for instance, is a well-known, effective camouflage strategy where the color and pattern of a stationary animal match its surrounding background. However, background matching may lose its efficacy when the animal moves, as the boundaries of the animal become more defined against its background. Recent evidence shows otherwise, as camouflaged objects can be less detectable than uncamouflaged objects even while in motion. Here, we explored if the detectability of computer-generated stimuli varies with the speed of motion, background (matching and unmatching) and size of stimuli in six species of jumping spiders (Araneae: Salticidae). Our results showed that in general, the responsiveness of all six salticid species tested decreased with increasing stimulus speed regardless of whether the stimuli were conspicuousness or camouflaged. Importantly, salticid responses to camouflaged stimuli were significantly lower compared to conspicuous stimuli. There were significant differences in motion detectability across species when the stimuli were conspicuous, suggesting differences in visual acuity in closely related species of jumping spiders. Furthermore, small stimuli elicited significantly lower responses than large stimuli across species and speeds. Our results thus suggest that background matching is effective even when stimuli are in motion, reducing the detectability of moving stimuli.

**Summary Statement:** Contrary to belief, stimuli in motion can remain camouflaged against their backgrounds. Using computer-generated stimuli, we show that smaller and faster stimuli against camouflaged background elicit lower responses from jumping spiders.

## INTRODUCTION

One of the most commonly known anti-predator strategies, background matching refers to the matching of an individual’s color and pattern to its background, thus rendering the individual to be indistinguishable from its surroundings (Michalis et al., 2017; Ruxton et al., 2019; Stevens and Merilaita, 2011). If the individual is successfully matched to its background, the probability of detection by visually oriented predators can be effectively lowered (Cuthill, 2019; Michalis et al., 2017). However, there may be limitations associated with this strategy. The benefits associated with matching a particular background type may be traded off against a different background type (Cuthill, 2019). More importantly, the edges of background matching individuals could become more noticeable against a patterned background when they move (Hall et al., 2013). This has thus led to the long-held belief that motion and camouflage are mutually exclusive (Cicerone and Hoffman, 1997; Ioannou and Krause, 2009; Regan and Beverley, 1984; Rushton et al., 2007).

Recent studies lend support to the notion that motion does not necessarily “break” camouflage, as there are various adaptations an individual can rely on to prevent detection whilst moving (Brunyé et al., 2019; Hall et al., 2013; Smart et al., 2020; Umeton et al., 2017; Yin et al., 2015). For instance, the presence of distractors or similarly patterned objects can improve the camouflaging effectiveness of an object (Hall et al., 2013). Brunyé et al. (2019) highlighted the importance of speed and pattern contrast in influencing detectability of moving stimuli. Furthermore, fast-moving prey tend to be harder to capture against a heterogeneous background compared to a uniform background (Stevens et al., 2008). A moving animal can also avoid detection by exploiting the receiver’s visual constraints through a combination of color patterns and behavioral adaptations aimed at misleading predators (e.g., protean motion (Chance and Russell, 1959), motion dazzle (Scott-Samuel et al., 2011)) or concealing motion signals (e.g., flicker fusion camouflage (Umeton et al., 2019), motion masquerade (Bian et al., 2016)). Although there are limitations associated with cryptic strategies in concealing moving individuals, there is some evidence showing that camouflaged targets are harder to detect than uncamouflaged or conspicuous targets (Brunyé et al., 2019; Stevens et al., 2008).

Whether a moving prey can be detected ultimately depends on the visual capabilities of the predator species. Most studies investigating the effect of movement on camouflage have been conducted using humans as predator models (Brunyé et al., 2019; Hall et al., 2013; Murali and Kodandaramaiah, 2018; Stevens et al., 2008). However, studies based on human perception may inevitably differ from those of natural predator-prey systems, thus making it difficult to infer the camouflage efficacy of the moving prey. Furthermore, studies of camouflage and motion are often challenging, as it is difficult to determine the effects of motion on the eyes of potential predators. For instance, prey may be easily detectable to predators with high visual acuity even if the prey moves at high speeds. Many studies of predator-prey interactions focus on the perception of avian predators, which possess excellent vision (e.g., Hodos, 2012; Jones et al., 2007). However, the camouflaging efficacy of moving prey may be higher for other potential predators with lower visual acuity, such as insects and spiders (Harland et al., 2012).

With most studies focusing on only one focal predator species (Hämäläinen et al., 2015; Mizutani et al., 2003; Umeton et al., 2019; Watanabe and Yano, 2013), there is limited empirical evidence comparing the effects of movement on prey camouflage across closely related species of natural predators. While closely related species have similar eye morphologies and visual systems, they may differ in terms of visual acuity (Dobberfuhl et al., 2005). For instance, the fruit fly *Drosophila melanogaster* Meigen, and the killer fly *Coenosia attenuata* Stein have highly similar lenses with similar optical properties, yet *C. attenuata* has much higher levels of spatial acuity and information transfer compared to *D. melanogaster* (Gonzalez-Bellido et al., 2011). To better understand the perception and responses of natural predators, it is thus important to examine a diversity of species, even when closely related.

Jumping spiders (Araneae: Salticidae), the most diverse clade amongst spiders (http://wsc.nmbe.ch, accessed on 3 May 2023), are well known for their high-resolution vision, which plays a critical role in their elaborate courtship and predatory behavior (Harland et al., 2012; Land, 1985). Many salticid species, such as *Portia* spp. And *Phidippus* spp., have especially high visual acuity relative to their body size (Cerveira et al., 2021; Harland et al., 1999; Land, 1985). Yet, three lineages within the family Salticidae – Lyssomaninae Blackwall, Spartaeinae Wanless, and Salticinae Blackwall – are known to differ greatly in terms of eye design and vision-based predatory strategies (Harland et al., 2012; Maddison, 2015; Maddison and Hedin, 2003; Su et al., 2007). As the structures and functions of salticids’ eyes are relatively well-studied (De Agrò et al., 2021; Harland et al., 2012; Jakob et al., 2018; Land, 1985; Zurek et al., 2010), salticids are good models to understand how they respond to signals in motion.

In this study, we investigated the responses of six species of salticids – *Cosmophasis umbratica* Simon, *Menemerus bivittatus* (Dufour), *Phintella vittata* (C. L. Koch), *Portia labiata* (Thorell), *Siler semiglaucus* (Simon), and *Thiania bhamoensis* Thorell *–* to moving stimuli against different backgrounds. First, we determined the responsiveness of salticids to conspicuous, differently sized stimuli in motion (i.e., visual responsiveness assay). Next, we examined the salticid responses to camouflaged stimuli at different speeds (i.e., background matching assay). We hypothesised that across the species, background matching can effectively camouflage motion, such that salticid response to background matching stimuli is lower than the response to conspicuous stimuli. We also expect that salticid responses to motion stimuli depend on species due to their potentially varying visual acuity.

## MATERIALS AND METHODS

### Study system and salticid maintenance

In this study, we used six salticid species commonly found in Singapore – *C. umbratica* (Salticinae: Saltafresia: Chrysillini), *M. bivittatus* (Salticinae: Saltafresia: Chrysillini), *Ph. vittata* (Salticinae: Saltafresia: Chrysillini), *Po. labiata* (Spartaeinae: Spartaeina), *S. semiglaucus* (Salticinae: Saltafresia: Chrysillini), and *T. bhamoensis* (Salticinae: Saltafresia: Euophryini) (Fig. 1). Additionally, *Ph. vittata* individuals (*N* = 8, 4 females and 4 males) were collected from Ipoh, Malaysia. A total of 191 salticids were used: *C. umbratica* (*N* = 39, 20 females and 19 males), *M. bivittatus* (*N* = 30, 16 females and 14 males), *Ph. vittata* (*N* = 40, 19 females and 21 males), *Po. labiata* (*N* = 12, 8 females and 4 males), *S. semiglaucus* (*N* = 33, 15 females and 18 males), and *T. bhamoensis* (*N* = 37, 19 females and 18 males).

**Fig. 1.**
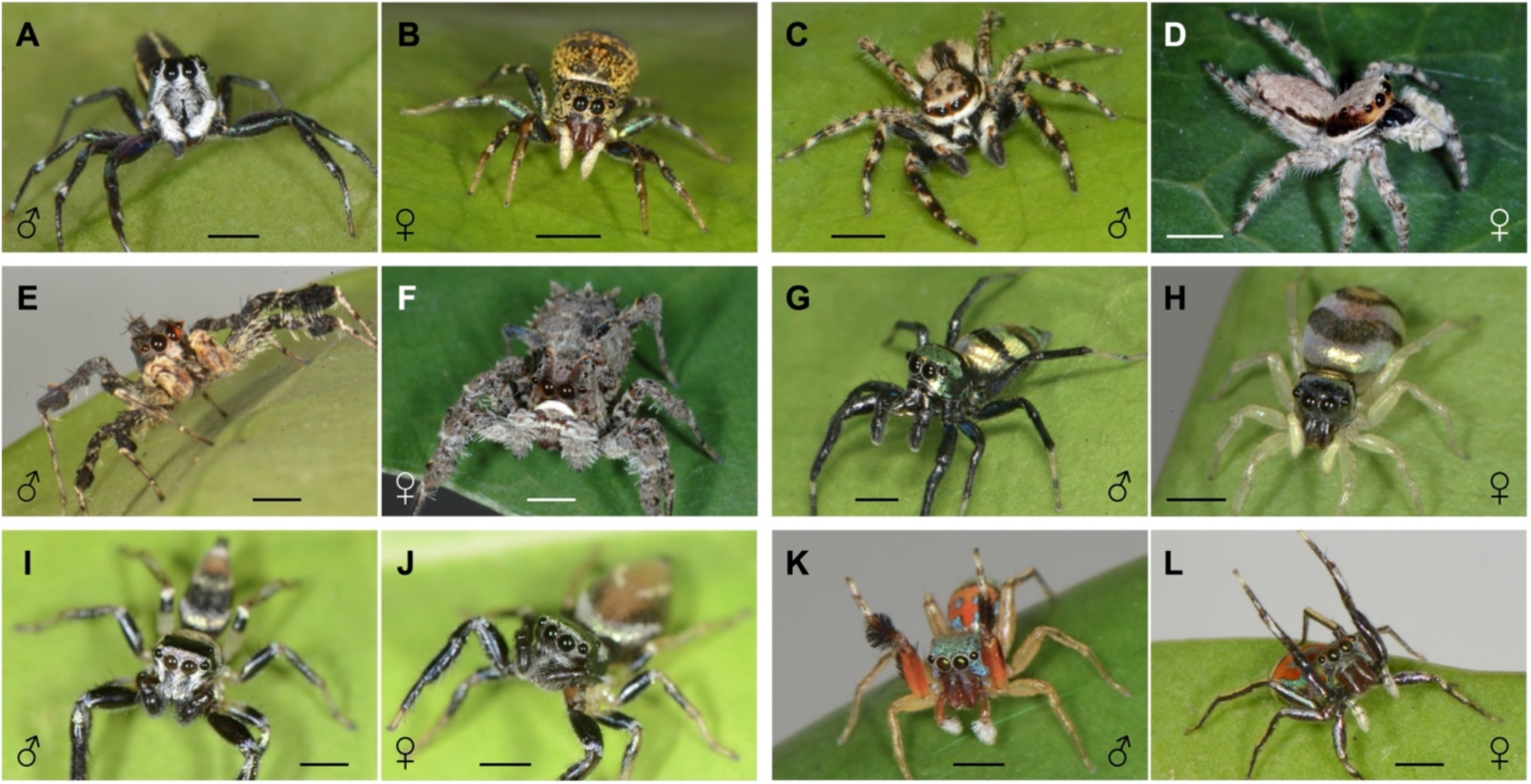
**Six species of salticids used in this study – *Cosmophasis umbratica* (A) male and (B) female; *Menemerus bivittatus* (C) male and (D) female; *Portia labiata* (E) male and (F) female; *Phintella vittata* (G) male and (H) female; *Thiania bhamoensis* (I) male and (J) female; and *Siler semiglaucus* (K) male and (L) female.**Each scale bar represents 2 mm in length approximately. Image credits: Min Tan, Daiqin Li, and Fan Li.

All salticids were housed individually in plastic containers (6 × 5 × 5 cm) under controlled environmental conditions (25 ± 1°C, 80 ± 5% RH, 12:12 h light: dark cycle). Each salticid was fed with five to seven laboratory-cultured *D. melanogaster* twice a week and provided with water *ad libitum*. As *Po. labiata* prefer spiders as prey (Harland and Jackson, 2000; Jackson and Hallas, 1986; Li et al., 1997) and were larger than the other species tested (pers. obs., Min Tan), they were fed small spiders once every two to three weeks. To simulate its natural habitat, we placed a small, dried leaf in the housing container for each individual *Po. labiata* for nest-building. When the experiments were completed, the salticids were euthanised using carbon dioxide and preserved in 75 % ethanol.

### General experimental procedures

To determine how the different salticid species respond to the stimuli moving at different speeds in the two assays, we used the experimental method adapted from Umeton et al. (2019), which involved presenting the test subject with computer generated stimuli moving at different speeds. We carried out two assays – the visual responsiveness assay and background matching assay – using a similar set-up (Fig. 2A) and procedures except that stimulus type and background were different between two assays (Fig. 2B, C; see details below). To prepare the background and the background matching stimuli for the background matching assay, we used images of the bark of rubber trees, *Hevea brasiliensis* Müll. Arg. The rubber tree can be widely found in Singapore, and several species of salticids (e.g., *M. bivittatus, Po. labiata*) and their prey are commonly found on the bark of rubber trees (pers. obs., Min Tan). Tree trunks were photographed at breast height (∼1.5 m), at a 0.5 m distance using a tripod-mounted Nikon D800 digital SLR camera (Nikon Corp., Tokyo, Japan). The aperture, shutter speed and ISO were kept at f16, 1/60 and 100 respectively. A total of 52 rubber tree trunks were imaged at Pulau Ubin, Singapore. Images were stored in RAW format to avoid loss of information due to compression (Stevens et al., 2007). We used the software ImageJ v.1.49 (Schneider et al., 2012) to calibrate and select a section of the bark photos. To prevent the tree’s curvature from distorting the measurement of the texture, only middle sections of the trunk (1200 × 2400 pixels) were used. We adjusted the brightness and contrast of the images before overlaying and stacking them into one image, thus creating a generic tree bark background. We then converted the background into a monochrome image and cropped the image to fit within the dimensions of a PowerPoint slide (25.4 × 19.05 cm, 960 × 720 pixels, 96 pixels per inch (ppi)).

**Fig. 2.**
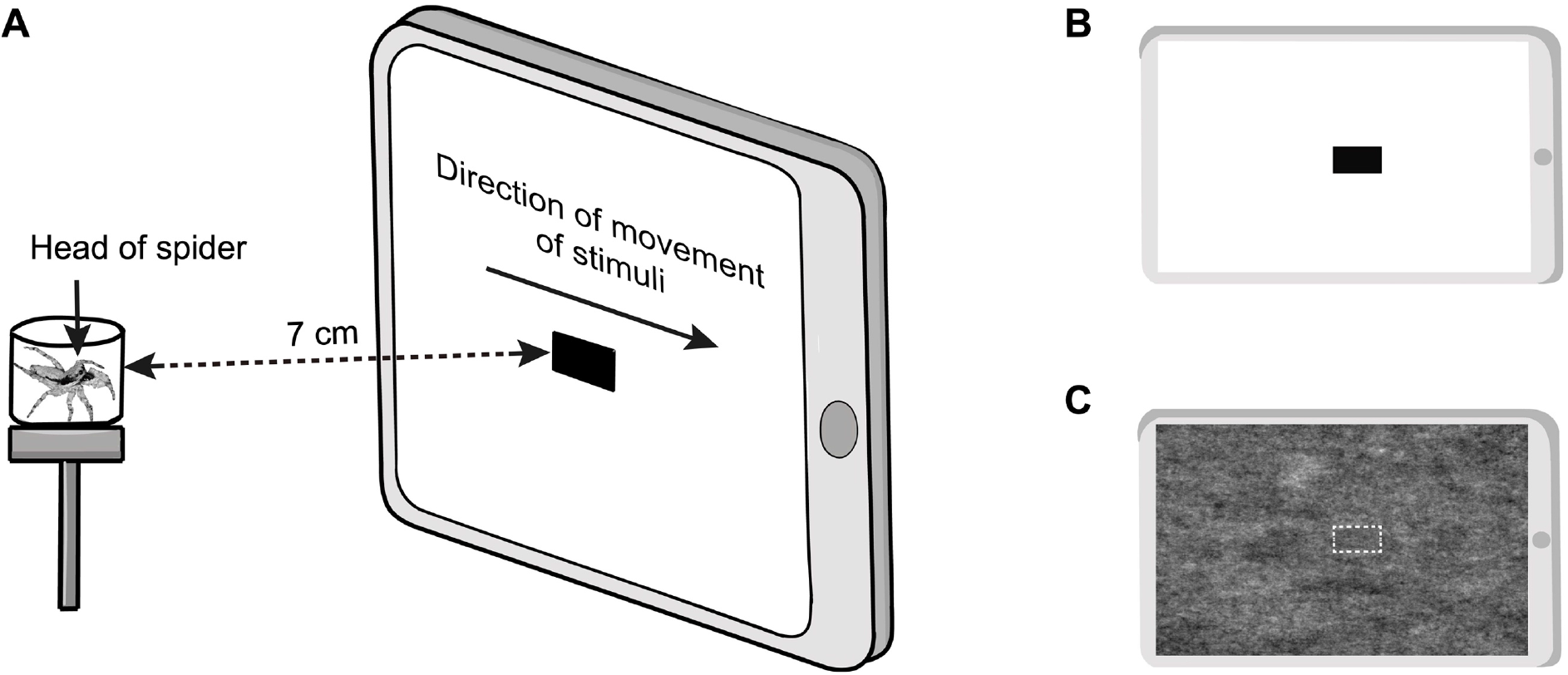
Experimental setup. (**A**) Stimuli were animated to move horizontally across the screen in front of the test salticid in a chamber, which is positioned at the centre of the tablet screen. (**B**) and **(C**) depict the stimuli and backgrounds for the visual responsiveness and background matching assays, respectively. The white border surrounding the background matching stimulus is only presented here for clarity and was not displayed in trials.

In both the visual responsiveness and background matching assays, the stimulus comprised of a rectangle moving horizontally across the screen. In the visual responsiveness assay, we used a black rectangle stimulus (< 1 cd m^-2^; Fig. 2B) presented against the white background. To test for the effects of size on visual responsiveness of motion, we used stimuli of two different sizes – large (63 × 30 pixels, corresponding to 1.68 × 0.79 cm, 96 ppi) and small (29 × 15 pixels, corresponding to 0.76 × 0.41 cm, 96 ppi). The large and small stimuli are comparable to the body lengths of a salticid and its potential prey, respectively. In the background matching assay, we used a background matching stimulus presented against the tree bark background (Fig. 2C). The background matching stimulus was extracted from the middle section of the background and the dimensions were the same as that used for the large stimulus in the visual responsiveness assay.

Before each trial, we placed the test salticid in a cylindrical, viewing chamber positioned on an elevated platform 7 cm from the centre of a tablet screen (iPad 8^th^ generation, Apple Inc, California, US), where the stimulus was displayed (Fig. 2A). Although salticids have been reported to discriminate prey from distances of up to 32 cm, some species, such as *Menemerus* sp., have a maximum discrimination distance of up to 12 cm (Harland et al., 1999). Thus, we deemed 7 cm to be an acceptable distance for our assays. The chamber consisted of a white base (diameter: 1.5 cm), a thin, transparent wall made of clear acetate (thickness: 0.1 cm, height: 0.7 cm) and a transparent lid (adhered to the wall using white tack). As *Po. labiata* individuals were generally larger than the rest of the species used, we placed them in a larger viewing chamber (diameter: 3 cm; height: 0.7 cm). The tablet screen was 20.8 × 15.6 cm, with a pixel resolution of 2160 × 1620 pixels (264 pixels per inch (ppi)) and a refresh rate of 60 Hz, which was deemed to be acceptable for this experiment as salticids can perceive flicker up to 40 Hz (Clark and Uetz, 1990). All stimulus animations were made using PowerPoint (Microsoft Corporation, New Mexico, US), which has a frame rate of 60 frames per second (fps). As salticids are generally diurnal (Foelix, 2011), all trials were conducted during daylight hours (i.e., 0900 – 1800 h). To ensure that the only light came from the iPad, we performed all trials in a dark room in absolute darkness. We recorded the trials using two video cameras (CASIO EX-100, Tokyo, Japan) positioned to record the anterior and posterior views of the test salticid.

The test salticid was first acclimatised in the chamber for 10 min. Next, we waved a thin brush in front of the test salticid to redirect its attention to the screen or gently rotated the chamber using a brush such that the salticid faced the screen. When the salticid turned towards the screen, the stimulus (Fig. 2B, C) was animated to move horizontally across the screen, with the direction of movement of stimuli randomised for each trial. The test ended after the stimulus animation moved across the screen. The stimuli and background varied depending on the assay.

### Visual responsiveness assay

To determine how salticids respond to an uncamouflaged stimulus, each test salticid was exposed to a black stimulus presented against the white background at different speeds. Each trial consisted of eight presentations, where the stimulus was presented in two sizes – large and small – at four moving speeds (duration of animation on screen: 0.25, 0.5, 1, and 1.5 s, corresponding to the speeds: high (83.2 cm s^-1^), medium (41.6 cm s^-1^), low (20.8 cm s^-1^), and very low (13.9 cm s^-1^)). The mean luminance of the white screen was approximately 300 cd/m^2^, measured using an illumination meter (Topcon IM-2D, Tokyo, Japan) at the start of each trial. Each presentation was separated by an inter-stimulus black screen interval of at least 50 s, and the stimuli were presented in a random order. During the assay, we observed that, unlike the other species tested, the behavior of *Po. labiata* was affected by the transition from a black to white screen, as *Po. labiata* would exhibit freezing behavior for a long time upon the screen transition. Thus, for *Po. labiata*, we used a white screen instead of a black screen during the inter-stimulus interval to minimise potential distractions for this species.

Each trial was repeated four times, with at least a one-day interval between each trial. Thus, a total of 32 stimulus presentations were showed to each salticid – each stimulus type (with two different sizes) presented at four speeds for four times.

### Background matching assay

To determine how salticids respond to a camouflaged stimulus moving at different speeds, salticids were tested using the similar procedures as in the visual responsiveness assay except that only the large, background matching stimuli (Fig. 2C) were presented against the tree bark background in the background matching assay. Thus, only essential details are described here. To ensure that the salticids were sufficiently responsive to the camouflaged stimulus, only individuals with an average response rate of at least 50 % to large, black stimuli in the visual responsiveness assay were included in this assay (Fig. S1). *Portia labiata* had very low response rate (only 25 % of *Po. labiata* had a response rate of at least 50 %) and were thus excluded from this assay. The mean luminance of the screen when the tree bark background was displayed was 45 cd m^-2^, measured using an illumination meter at the start of each trial. Each salticid was exposed to a trial consisting of four stimuli presentations, where the background matching stimulus was presented at four speeds (corresponding to the speeds tested in the visual responsiveness assay) in a random order. We repeated each experimental trial four times, with at least a one-day interval between each trial. Thus, a total of 16 stimuli were presented to each salticid.

### Behavioral responses

To determine whether the stimulus was detected by the salticids, we scored the responses of the test salticids following each stimulus presentation (Table 1; Movie 1).

**Table 1.** Behavioral responses observed during the assays.

| Behavioral response type | Description |
| --- | --- |
| No response | No observable change in behavior. |
| Mild response | Pausing or freezing behavior including paused movement of pedipalps.<br>Shifting of legs or abdomen.<br>Delayed response after the stimulus passed the screen. |
| Strong response | Turning of head in response to the stimulus.<br>Moving towards, retreating away, or flinching from the stimulus. |

### Statistical analysis

We performed two analyses to determine – i) the effectiveness of background matching for moving stimuli; ii) if there are variations in responses among salticid species. For both analyses, we proposed a series of cumulative link mixed models (CLMM) using the *ordinal* package (https://CRAN.R-project.org/package=ordinal). We coded the behavioral responses of salticids in three levels as the response variable. Salticid identity (ID) and trial number (i.e., sequence of trial for each salticid) were coded as the random intercept and slope, respectively, for all CLMM. We then ranked these models using Akaike’s information criterion for small sample sizes (AICc; Burnham and Anderson, 2002) and identified the best-fitting model using *model*.*sel* in the *MuMIn* package (http://R-Forge.R-project.org/projects/mumin). We performed all analyses using R version 4.3.0 (http://www.R-project.org/).

To determine the effectiveness of background matching for moving stimuli, we compared the salticids’ responses across both assays and included salticid species, stimulus moving speed, salticid sex, and background type as fixed effects. We proposed a total of 14 models, which comprised of i) a null model (1 model); ii) each fixed effect alone (4 models); iii) salticid species interacting with the other fixed effects (3 models); iv) stimulus speed interacting with background/stimulus type (1 model); v) full models containing all fixed effects with and without interactions (5 models). A comparison of the models can be found in Table A1.

To determine if there are variations in responses among salticid species, we examined the salticids’ responses in the visual responsiveness assay only. We included the same fixed effects and models as above but replaced background type with stimulus size. The proposed models are ranked in Table A2. Additionally, we determined if there were differences in responses among species at each of the four stimulus speeds tested by using four separate CLMMs to compare the responses among species at each speed in the visual responsiveness assay. For each CLMM, salticid species, stimulus size and sex were included as fixed effects. We then conducted pairwise comparison tests for significant effects using *emmeans* (https://CRAN.R-project.org/package=emmeans) for each CLMM.

## RESULTS

### The effectiveness of background matching

The model containing species, stimulus moving speed, sex, background type, as well as the interaction between species and stimulus moving speed best predicted the responses of salticids between camouflaged and uncamouflaged stimuli (AICc = 3971.4, weight = 1; Table A1). Salticids generally showed a significantly higher response to uncamouflaged stimuli when compared to camouflaged stimuli regardless of species (Table 2; Fig. 3), indicating that background matching effectively reduced the detectability of moving stimuli for salticids.

**Table 2.** The best-fitting cumulative link mixed model showing the effects of species, sex, stimulus moving speed and background type on the level of responsiveness of salticids (i.e., no response, mild, and strong response).

| Predictors | Estimate | SE | Z | P |
| --- | --- | --- | --- | --- |
| **Background Type (camouflaged)** | **-1.1086** | **0.0900** | **-12.317** | **<0.0001** |
| Sex (male) | 0.0304 | 0.1141 | 0.266 | 0.790 |
| **Stimulus Speed** | **-0.0199** | **0.0036** | **-5.460** | **<0.0001** |
| *Menemerus bivittatus* | 0.3895 | 0.2509 | 1.553 | 0.1205 |
| ***Phintella vittata*** | **0.8271** | **0.2629** | **3.146** | **<0.01** |
| *Siler semiglaucus* | 0.1993 | 0.2766 | 0.721 | 0.4712 |
| ***Thiania bhamoensis*** | **1.1456** | **0.2769** | **4.137** | **<0.0001** |
| ***Menemerus bivittatus*: Stimulus Speed** | **-0.0111** | **0.0052** | **-2.153** | **0.0313** |
| ***Phintella vittata*: Stimulus Speed** | **-0.0158** | **0.0054** | **-2.953** | **<0.01** |
| *Siler semiglaucus*: Stimulus Speed | -0.0035 | 0.0055 | -0.634 | 0.5261 |
| ***Thiania bhamoensis*: Stimulus Speed** | **-0.0260** | **0.0059** | **-4.434** | **<0.0001** |

Significant effects are in bold. Trial number and salticid ID were assigned as random effects. *Cosmophasis umbratica*, background type (uncamouflaged) and females are used as references for species, stimulus size and sex, respectively.

**Fig. 3.**
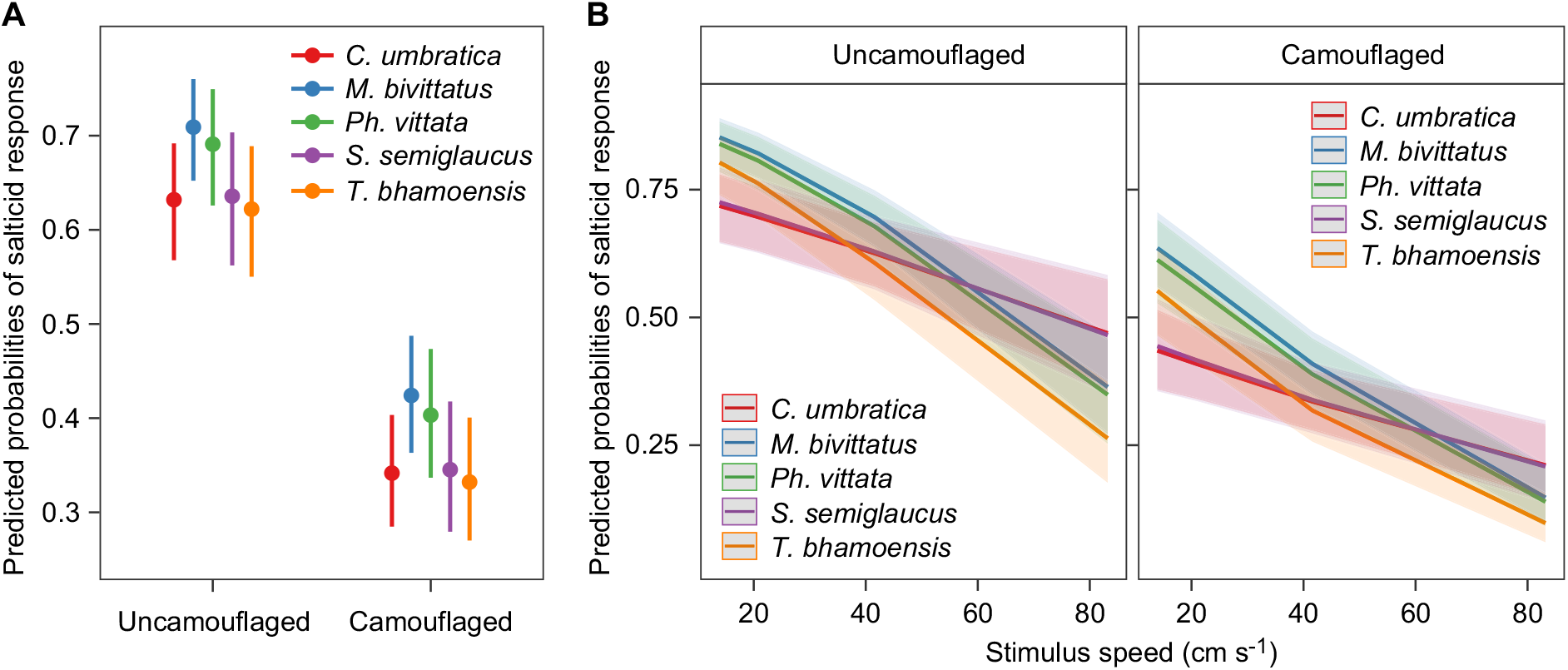
Predicted probabilities of the salticids’ responses (n=70) to camouflaged and uncamouflaged stimuli across five species (A) and stimulus moving speed (B). The error bars and shaded regions in **A** and **B**, respectively, represent 95 % c.i.

Salticids responded less with increasing stimulus moving speed across species, sexes, and backgrounds; salticids generally responded differently among species across stimulus moving speeds, sexes and backgrounds. However, females had no significant differences in responsiveness compared to males across species, stimulus moving speeds and backgrounds (Table 2).

### Differences in motion detectability among species

In the visual responsiveness assay, where the black stimuli were presented against the white background, the best-fitting model predicting salticid responses includes species, stimulus moving speed, stimulus size, sex, as well as the interaction between species and stimulus moving speed (AICc = 9872.6, weight = 1, Table A2). In general, the conspicuous stimuli elicited different levels of responses among the salticids species tested (Fig. 4A). Salticids responded significantly less with increasing moving speed, and small stimuli elicited significantly lower responses than large stimuli (Table 3 and Fig. 4B). However, sex did not significantly predict salticid responses though it is included in the best-fitting model (Table 3). Species interacting with stimulus moving speed significantly predicted salticid responses to conspicuous, moving stimuli: salticids that responded differently to the different stimulus moving speeds depended on salticid species (Table 3).

**Table 3.**
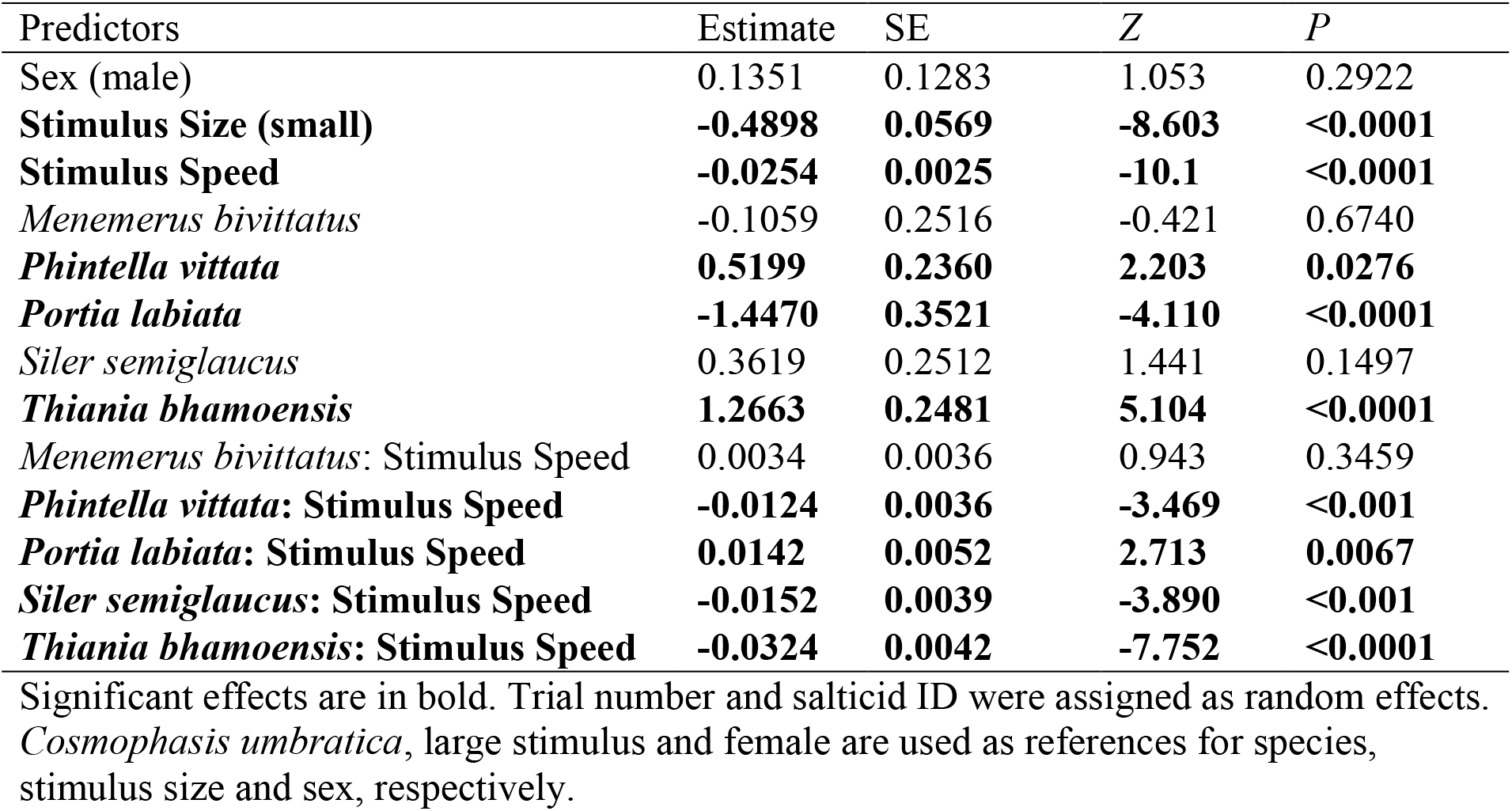
The best-fitting cumulative link mixed model reporting the effects of species, sex, stimulus moving speed, and size and the interaction between species and stimulus moving speed on the level of responsiveness of salticids (i.e., no response, mild, and strong response) in the visual responsiveness assay.

| Predictors | Estimate | SE | Z | P |
| --- | --- | --- | --- | --- |
| Sex (male) | 0.1351 | 0.1283 | 1.053 | 0.2922 |
| **Stimulus Size (small)** | **-0.4898** | **0.0569** | **-8.603** | **<0.0001** |
| **Stimulus Speed** | **-0.0254** | **0.0025** | **-10.1** | **<0.0001** |
| *Menemerus bivittatus* | -0.1059 | 0.2516 | -0.421 | 0.6740 |
| ***Phintella vittata*** | **0.5199** | **0.2360** | **2.203** | **0.0276** |
| ***Portia labiata*** | **-1.4470** | **0.3521** | **-4.110** | **<0.0001** |
| *Siler semiglaucus* | 0.3619 | 0.2512 | 1.441 | 0.1497 |
| ***Thiania bhamoensis*** | **1.2663** | **0.2481** | **5.104** | **<0.0001** |
| *Menemerus bivittatus*: Stimulus Speed | 0.0034 | 0.0036 | 0.943 | 0.3459 |
| ***Phintella vittata*: Stimulus Speed** | **-0.0124** | **0.0036** | **-3.469** | **<0.001** |
| ***Portia labiata*: Stimulus Speed** | **0.0142** | **0.0052** | **2.713** | **0.0067** |
| ***Siler semiglaucus*: Stimulus Speed** | **-0.0152** | **0.0039** | **-3.890** | **<0.001** |
| ***Thiania bhamoensis*: Stimulus Speed** | **-0.0324** | **0.0042** | **-7.752** | **<0.0001** |

Significant effects are in bold. Trial number and salticid ID were assigned as random effects. *Cosmophasis umbratica*, large stimulus and female are used as references for species, stimulus size and sex, respectively.

**Fig. 4.**
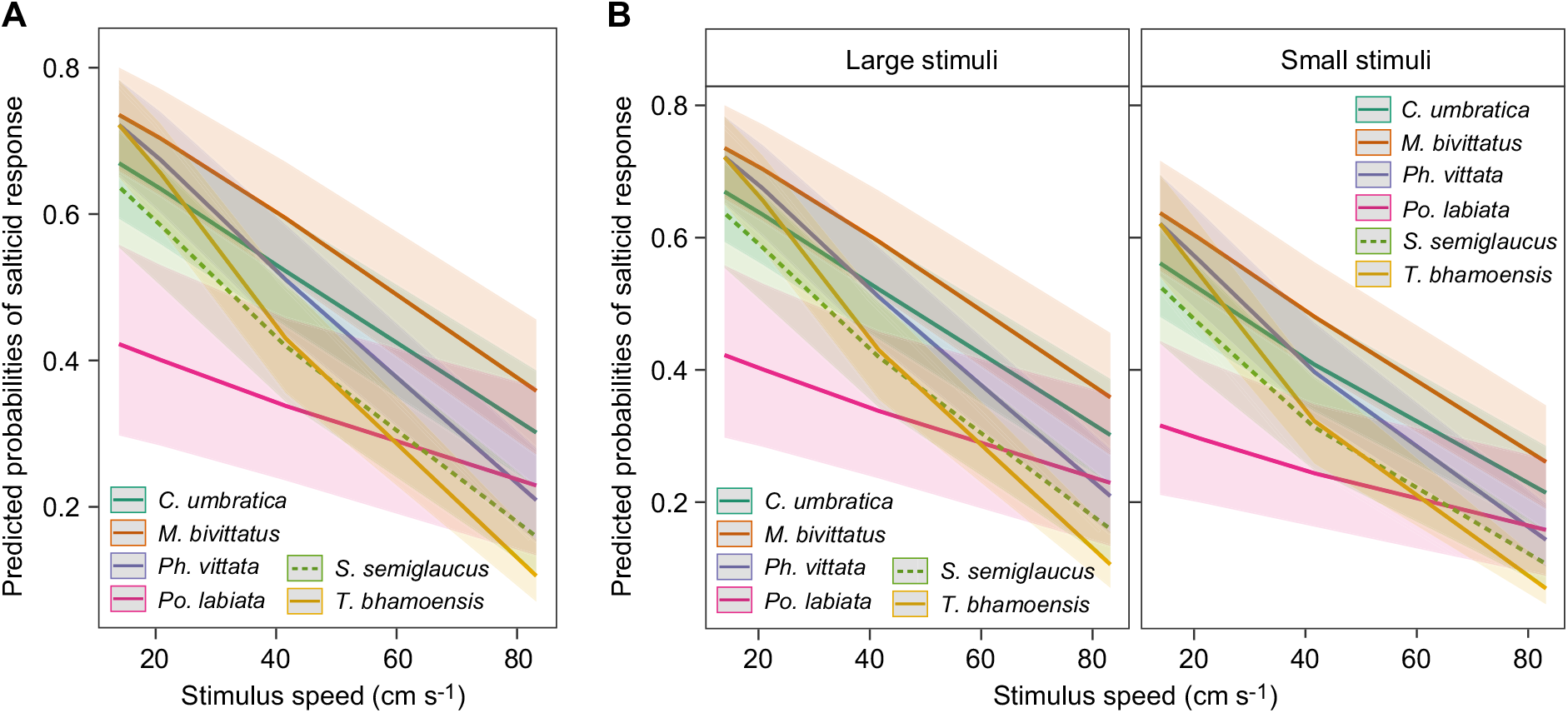
**Predicted probabilities of the salticids’ responses (n=191) (A) across the tested stimulus moving speed in the visual responsiveness assay and (B) to large (1.68 × 0.79 cm) and small (0.76 × 0.41 cm) stimuli.** Shaded regions represent 95 % c.i.

Pairwise comparisons of the salticids’ level of responsiveness across averaged levels of speed, size and sex in the visual responsiveness assay showed that *M. bivittatus, Ph. vittata*, and *T. bhamoensis* displayed significantly stronger responses compared to *Po. labiata* (Table 4). Furthermore, analyses at specific speeds showed significant differences in the salticids’ level of responsiveness (Fig. 4). *Portia labiata* was less responsive, and displayed milder responses compared to the rest of the species at most stimulus moving speeds (Fig. 5). Interestingly, *T. bhamoensis* displayed stronger responses compared to the rest of the species at lower speeds (Fig. 5). At the highest speed, *C. umbratica* and *M. bivittatus* displayed significantly stronger responses than *T. bhamoensis* (Fig. 5). No significant difference in level of responsiveness was found at the medium speed (Fig. 5).

**Table 4.** Pairwise comparisons of the level of response (i.e., no response, mild, and strong response) among all six species over the average of speed, size and sex in the visual responsiveness assay.

| Contrast | Estimate | SE | Z | P |
| --- | --- | --- | --- | --- |
| *Cosmophasis umbratica* – *Menemerus bivittatus* | -0.0293 | 0.214 | -0.137 | 1.0000 |
| *Cosmophasis umbratica* – *Phintella vittata* | -0.0228 | 0.199 | -0.114 | 1.0000 |
| ***Cosmophasis umbratica* – *Portia labiata*** | **0.8806** | **0.296** | **2.971** | **0.0352** |
| *Cosmophasis umbratica* – *Siler semiglaucus* | 0.2461 | 0.212 | 1.160 | 0.8557 |
| *Cosmophasis umbratica* – *Thiania bhamoensis* | 0.0272 | 0.206 | 0.132 | 1.0000 |
| *Menemerus bivittatus* – *Phintella vittata* | 0.0065 | 0.213 | 0.031 | 1.0000 |
| *Menemerus bivittatus* – *Portia labiata* | **0.9100** | **0.304** | **2.995** | **0.0328** |
| *Menemerus bivittatus* – *Siler semiglaucus* | 0.2754 | 0.221 | 1.244 | 0.8151 |
| *Menemerus bivittatus* – *Thiania bhamoensis* | 0.0564 | 0.218 | 0.259 | 0.9998 |
| *Phintella vittata* – *Portia labiata* | **0.9034** | **0.296** | **3.055** | **0.0273** |
| *Phintella vittata* – *Siler semiglaucus* | 0.2689 | 0.210 | 1.278 | 0.7971 |
| *Phintella vittata* – *Thiania bhamoensis* | 0.0500 | 0.205 | 0.244 | 0.9999 |
| *Portia labiata* – *Siler semiglaucus* | -0.6345 | 0.303 | -2.093 | 0.2907 |
| *Portia labiata* – *Thiania bhamoensis* | **-0.8534** | **0.300** | **-2.849** | **0.0501** |
| *Siler semiglaucus* – *Thiania bhamoensis* | -0.2189 | 0.216 | -1.015 | 0.9131 |

Significant effects are in bold.

**Fig. 5.**
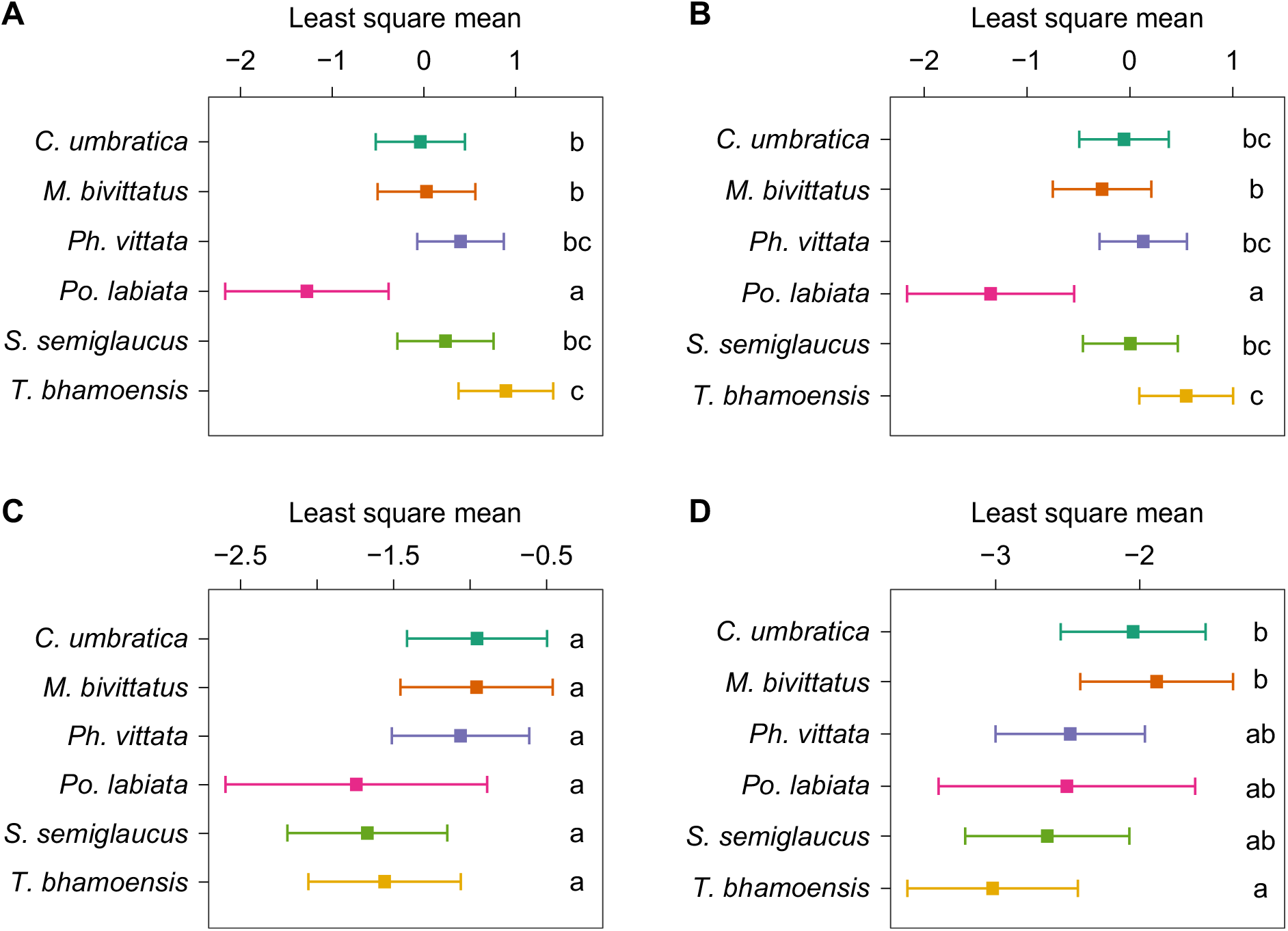
**Least squares (LS) mean for the six salticid species (n=191) – *Cosmophasis umbratica, Menemerus bivittatus, Phintella vittata, Portia labiata, Siler semiglaucus*, and *Thiania bhamoensis* – at each speed in the visual responsiveness assay.** Speeds: (**A**) Very low (13.9 cm s^-1^); (**B**) low (20.8 cm s^-1^); (**C**) medium (41.6 cm s^-1^); (**D**) high (83.2 cm s^-1^). Error bars indicate the 95 % c.i. of the LS mean. Different letters indicate Sidak-adjusted comparisons with significant differences.

## DISCUSSION

Our study demonstrates that salticids displayed lower levels of response to camouflaged than uncamouflaged stimuli, suggesting that background matching may be an effective camouflage strategy for moving stimuli, especially at higher speeds, in the eyes of salticids. Salticids displayed higher levels of response when the stimulus was larger and moving slower, indicating that the speed of moving stimuli inversely correspond to detectability.

Importantly, we observed significant differences in motion detectability across species when the stimuli were conspicuous, thus suggesting differences in visual acuity across closely related salticid species.

Our findings support our hypothesis that camouflaged moving stimuli, compared to uncamouflaged moving stimuli, were less detectable across the salticid species tested in this study. One possible reason can be that the optical performance of these salticid species tested may not be well equipped in discriminating stimuli moving against a similarly patterned background. Although little is known regarding the visual acuity of the tested species, our results suggest that the salticids may have difficulty resolving complex textures such as the bark of rubber trees. Understanding the effectiveness of background matching for camouflage can provide insights to how moving animals exploit the visual constraints of the receiver to escape detection and capture. This is relevant as animals need to move to forage, find mates or escape from potential predators. Previous studies indicate that when a background matching prey moves, its boundaries become more defined, rendering them more detectable to predators (Hall et al., 2013; Ioannou and Krause, 2009). However, recent empirical studies suggest that prey detectability depends on the background – background matching prey can be more difficult to detect compared to uncamouflaged prey when coupled with a patterned background (Brunyé et al., 2019; Stevens et al., 2008). Perhaps, prey in motion may be more easily detected if the predator is actively searching for the prey. Unless it is already spotted, prey moving in the predator’s peripheral vision can escape detection (Smart et al., 2020).

Thus, a well-camouflaged prey would be able to escape detection more easily than a conspicuous one. Although our results indicate that background matching stimuli in motion experience reduced detection, it remains to be shown if these stimuli evade capture more successfully than non-background matching stimuli when detected. In addition, the effect of different motion types (e.g., irregular bursts of speed or unpredictable motion trajectory) could further improve the camouflaging effectiveness of background matching in moving prey.

As predicted, the responses of the salticids were negatively correlated with the speed of moving stimuli regardless of species, sex, and background type. Our findings are in line with prior studies, which found that prey moving at higher speeds tend to avoid capture (Smart et al., 2020; Stevens et al., 2008). At high speeds, the stimuli could have moved so quickly that it exceeded the motion processing capabilities of the salticids, thus resulting in lower response levels. Additionally, fast moving stimuli have an added advantage in avoiding detection if it is moving against a similarly patterned background. Thus, in our study, camouflaged stimuli elicited lower responses compared to conspicuous stimuli, especially at high speeds.

The variation in responses to moving stimuli among salticid species may be explained by their phylogenetic relationship. Five out of the six species tested in this study belong to the subfamily Salticinae – *C. umbratica, M. bivittatus Ph. vittata, S. semiglaucus* and *T. bhamoensis*, while *Po. labiata* belongs to the subfamily Spartaeinae (Maddison, 2015). While the five species belong to the same clade Saltafresia Bodner & Maddison, *T. bhamoensis* belongs to the tribe Euophryini Simon, whereas the other four species belong to the tribe Chrysillini Simon (Maddison, 2015). Although species of the subfamily Salticinae are known to have high spatial acuity (Cerveira et al., 2021; Harland et al., 2012; Su et al., 2007), much remains unknown regarding differences in eye structure and visual acuity between Euophryini and Chrysillini. At low speeds, *T. bhamoensis* displayed stronger responses compared to the rest of the species. Conversely, we observed stronger responses for *C. umbratica* and *M. bivittatus* at high speeds. Our results suggest that Euophryini and Chrysillini salticids have variations in motion responsiveness depending on the speed – Euophryini and Chrysillini salticids seem to have higher visual acuity at low and high speeds, respectively.

Variation in responses across species may be the result of adaptations to different habitats (Cerveira et al., 2021; Steinhoff et al., 2020). The salticid species tested in this study were found in habitats with different levels of ambient light. *Cosmophasis umbratica, Ph. vittata, S. semiglaucus*, and *T. bhamoensis* are typically found in garden bushes and shrubs (Koh et al., 2022). On the other hand, *M. bivittatus* and *Po. labiata* are more commonly found on tree trunks instead (Cerveira et al., 2021; Maddison, 2015). *Menemerus bivittatus* can also be found on human infrastructure, such as walls and railings, (pers. obs., Min Tan), which can vary greatly in terms of lighting. Luminance is an important factor in motion perception since it is necessary in the discrimination of moving objects (Troscianko et al., 2009). Thus,

*M. bivittatus* may be more accustomed to movement in dynamic environmental conditions compared to the other species tested in our study. Variations in luminosity could have also influenced stimulus detectability against different background types. In our study, the textured background had lower mean luminosity compared to the more conspicuous white background. Thus, it is possible that the camouflaged stimuli appear to be less conspicuous than the uncamouflaged stimuli due to differences in luminosity.

Low levels of response may not represent low motion processing capabilities but could be due to species-specific responses. *Portia labiata* displayed low levels of responses compared to the other species of salticids. However, *Portia* species are known to have high spatial acuity and greater resolving power compared to other salticids (Harland et al., 2012; Land, 1985). Spiders of the genus *Portia* behave differently from the majority of salticids, such as adopting a “choppy” and slow gait (i.e., jerking their legs and pedipalps and pausing at irregular intervals) to better camouflage them against their surroundings (Jackson and Blest, 1982), and preferentially practice araneophagy, where they have evolved specialised and complex strategies to capture arachnid prey (Harland and Jackson, 2000). When faced with a potential salticid prey, *Portia* spp. tend to freeze until the prey turned away (Jackson and Blest, 1982). The low level of response of *Po. labiata* in our study can therefore be likely attributed to their prey-stalking strategy, rather than low visual acuity and motion detectability capabilities.

Larger stimuli may elicit greater response than small stimuli through different mechanisms of perception biases and optimal foraging. When moving, the size of an individual prey can alter the viewer’s depth perception. For instance, larger objects may be perceived closer to the viewer compared to a smaller object (Scott-Samuel et al., 2011).

Furthermore, large targets can appear to move more slowly than small targets (Brown, 1931). Prey body size is shown to influence a predator’s foraging decision, especially since the predator needs to evaluate if the rewards obtained from prey capture outweigh the costs/ risks involved (Juanes, 1992; Nakazawa et al., 2013). Oftentimes, predators must balance potential risks and rewards before pursing prey – a relatively large prey may be energetically costly to predators while a small prey may not yield sufficient energetic returns (Schmitz, 2017). In our study, salticids might have been more responsive to large stimuli as larger stimuli would return greater rewards. Alternatively, since the large stimuli is comparable to the size of a salticid, the salticids may judge the large stimulus to be a greater threat than small stimulus and, thus, displayed heightened levels of caution by showing stronger response.

Our findings highlight the importance of stimuli speed and size in camouflaging motion, and how the receiver’s responses can vary in motion perception across salticid species within the same subfamily and tribe. We provide empirical support on how background matching aids the concealment of moving stimuli. Future studies can further build on these findings to better understand the ecological relevance of camouflage strategies in relation to motion.

## Acknowledgements

We would like to thank Boon Hui Wong, Tammy Ai Tian Ho, Bernetta Zi Wei Kwek, and Rachel Wan Xin Seah for assistance in spider collection, and Lu Wee Tan for managing the insectary.

## Author contributions

Conceptualization: M.T., D.L., E.J.T.; Methodology: M.T., D.L., E.J.T., J.Y.O.C.; Formal analysis: M.T., D.L., E.J.T.; Investigation: M.T., D.L., E.J.T., J.Y.O.C.; Resources: M.T., D.L., E.J.T.; Writing - original draft: M.T.; Writing - review & editing: M.T., D.L., E.J.T., J.Y.O.C., L.Y.; Visualization: M.T., L.Y.; Supervision: D.L., E.J.T.; Project administration: D.L., E.J.T.; Funding acquisition: D.L., E.J.T.

## Data availability

Data, analyses and codes in this article are available from the Dryad Digital Repository: https://datadryad.org/stash/share/mIX8RUcboFTT8woRK70eSuleqUBkPLGrscfOvNAvTc8

## APPENDICES

**Table A1.** Comparison of 14 CLMM to determine the effectiveness of salticid species, sex, stimulus speed and background type on the salticids’ response level (i.e., no response, mild response, and strong response).

| Predictors | df | logLik | AICc | delta | weight |
| --- | --- | --- | --- | --- | --- |
| Background Type + Sex + Stimulus Speed * Species | 16 | -1969.5 | 3971.2 | 0 | 1 |
| Sex + Stimulus Speed * Background Type + Species | 13 | -1982.1 | 3990.4 | 19.2 | 0 |
| Stimulus Speed * Background Type | 8 | -2023.7 | 4063.5 | 92.3 | 0 |
| Sex + Stimulus Speed + Background Type * Species | 16 | -2022.0 | 4076.3 | 105.1 | 0 |
| Background Type + Sex + Stimulus Speed + Species | 12 | -2026.2 | 4076.6 | 105.4 | 0 |
| Background Type + Stimulus Speed + Sex * Species | 16 | -2030.6 | 4093.4 | 122.2 | 0 |
| Species * Stimulus Speed | 14 | -2058.3 | 4144.8 | 173.6 | 0 |
| Stimulus Speed | 6 | -2081.4 | 4174.7 | 203.5 | 0 |
| Background Type | 6 | -2149.1 | 4310.2 | 339.0 | 0 |
| Species * Background Type | 14 | -2143.2 | 4314.7 | 343.5 | 0 |
| Null | 5 | -2215.0 | 4440.1 | 468.9 | 0 |
| Sex | 6 | -2215.0 | 4442 | 470.8 | 0 |
| Species | 9 | -2213.4 | 4444.9 | 473.7 | 0 |
| Species * Sex | 14 | -2212.5 | 4453.2 | 482.0 | 0 |

**Table A2.** Comparison of 14 CLMM to determine the effectiveness of salticid species, sex, stimulus speed and size on the salticids’ response level (i.e., no response, mild response, and strong response).

| Predictors | df | logLik | AICc | delta | weight |
| --- | --- | --- | --- | --- | --- |
| Sex + Stimulus Size + Stimulus Speed*Species | 18 | -4916.5 | 9869.2 | 0 | 1 |
| Species * Stimulus Speed | 16 | -4954.7 | 9941.5 | 72.3 | 0 |
| Sex + Stimulus Size + Stimulus Speed + Species | 13 | -4978.5 | 9983 | 113.8 | 0 |
| Sex + Stimulus Speed + Stimulus Size * Species | 18 | -4976.7 | 9989.6 | 120.4 | 0 |
| Stimulus Speed * Stimulus Size | 8 | -4987.4 | 9990.7 | 121.5 | 0 |
| Sex * Species + Stimulus Speed + Stimulus Size | 18 | -4984.0 | 10004.2 | 135 | 0 |
| Sex + Stimulus Size * Stimulus Speed + Species | 14 | -5070.7 | 10169.4 | 300.2 | 0 |
| Stimulus Speed | 6 | -5128.6 | 10269.2 | 400.0 | 0 |
| Species * Stimulus Size | 16 | -5453.9 | 10939.9 | 1070.7 | 0 |
| Stimulus Size | 6 | -5464.8 | 10941.6 | 1072.4 | 0 |
| Species | 10 | -5483.7 | 10987.5 | 1118.3 | 0 |
| Species * Sex | 16 | -5481.4 | 10995 | 1125.8 | 0 |
| Null | 5 | -5492.9 | 10995.9 | 1126.7 | 0 |
| Sex | 6 | -5492.2 | 10996.5 | 1127.3 | 0 |

### SUPPLEMENTARY MATERIALS

**Fig. S1.**
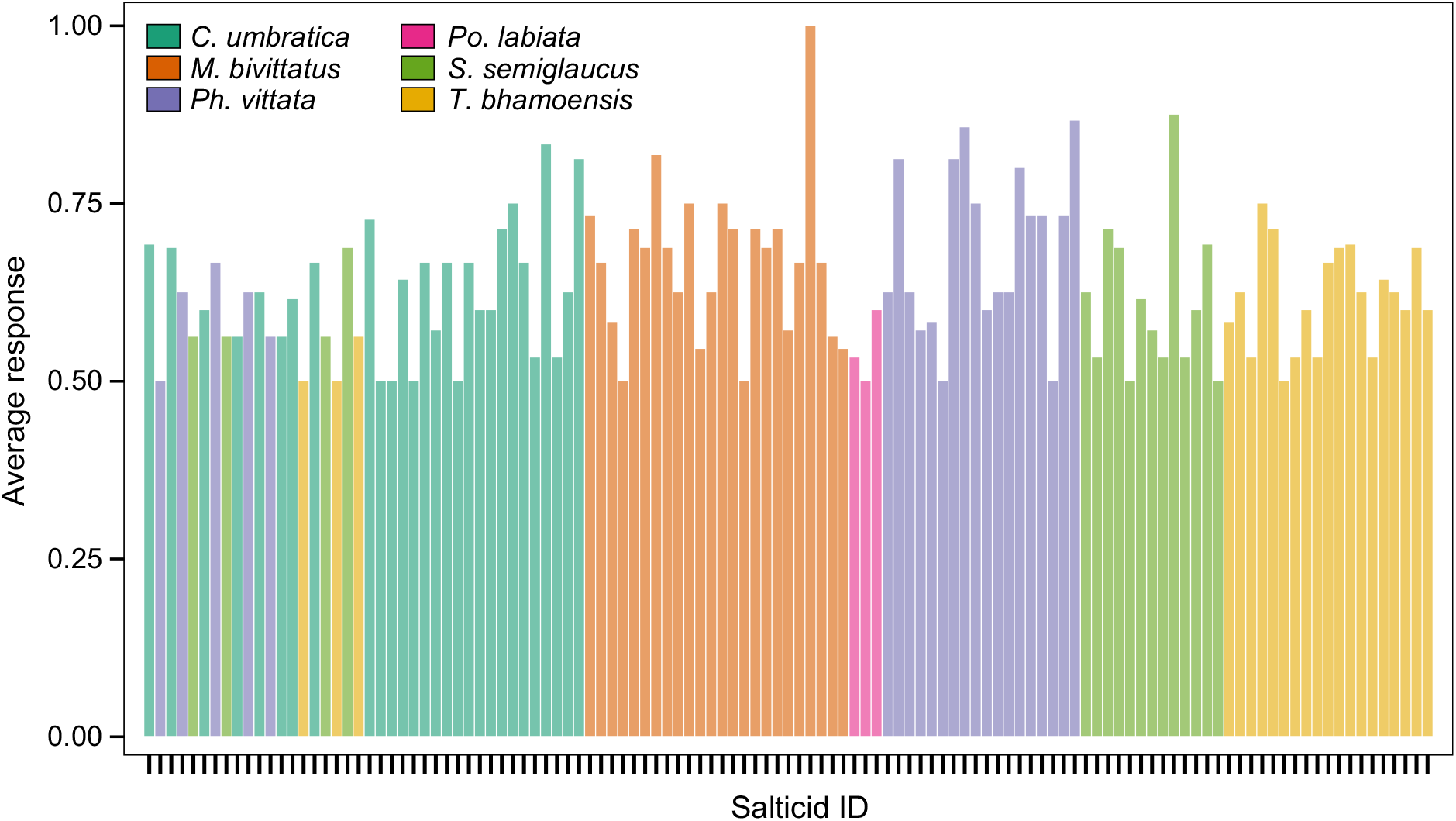
Salticids with an average response rate to large, black stimuli of at least 50 % in the visual responsiveness assay.

## References

Bian, X., Elgar, M. A. and Peters, R. A. (2016). The swaying behavior of Extatosoma tiaratum: motion camouflage in a stick insect? Behav. Ecol. 27, 83–92. doi:10.1093/beheco/arv125

Brown, J. F. (1931). The visual perception of velocity. Psychol. Forsch. 14, 199–232. doi: 10.1007/BF00403873

Brunyé, T. T., Martis, S. B., Kirejczyk, J. A. and Rock, K. (2019). Camouflage pattern features interact with movement speed to determine human target detectability. Appl Ergon. 77, 50–57. doi:10.1016/j.apergo.2019.01.004

Burnham, K. P. and Anderson, D. R. (2002). Model selection and multimodel inference 2: A Practical Information-Theoretic Approach. New York, US: Springer.

Cerveira, A. M., Nelson, X. J. and Jackson, R. R. (2021). Spatial acuity-sensitivity trade-off in the principal eyes of a jumping spider: possible adaptations to a ‘blended’ lifestyle. J. Comp. Physiol. A. 207, 437–448. doi:10.1007/s00359-021-01486-2

Chance, M. and Russell, W. (1959). Protean displays: a form of allaesthetic behaviour. In Proc. Zool. Soc. Lond, pp. 65–70. UK: Blackwell Publishing Ltd.

Cicerone, C. M. and Hoffman, D. D. (1997). Color from motion: dichoptic activation and a possible role in breaking camouflage. Perception. 26, 1367–1380. doi:10.1068/p261367

Clark, D. L. and Uetz, G. W. (1990). Video image recognition by the jumping spider, Maevia inclemens (Araneae: Salticidae). Anim. Behav. 40, 884–890. doi:10.1016/S0003-3472(05)80990-X

Cuthill, I. C. (2019). Camouflage. J. Zool. 308, 75–92. doi:10.1111/jzo.12682

De Agrò, M., Rößler, D. C., Kim, K. and Shamble, P. S. (2021). Perception of biological motion by jumping spiders. PLoS Biol. 19, e3001172. doi:10.1371/journal.pbio.3001172

Dobberfuhl, A. P., Ullmann, J. F. and Shumway, C. A. (2005). Visual acuity, environmental complexity, and social organization in African cichlid fishes. Behav. Neurosci. 119, 1648. doi:10.1037/0735-7044.119.6.1648

Foelix, R. (2011). Biology of Spiders. Oxford, UK: Oxford University Press.

Gonzalez-Bellido, P. T., Wardill, T. J. and Juusola, M. (2011). Compound eyes and retinal information processing in miniature dipteran species match their specific ecological demands. Proc. Natl. Acad. Sci. USA. 108, 4224–4229. doi:10.1073/pnas.101443810

Hall, J. R., Cuthill, I. C., Baddeley, R., Shohet, A. J. and Scott-Samuel, N. E. (2013). Camouflage, detection and identification of moving targets. Proc. R. Soc. B. 280, 20130064. doi:10.1098/rspb.2013.0064

Hämäläinen, L., Valkonen, J., Mappes, J. and Rojas, B. (2015). Visual illusions in predator–prey interactions: birds find moving patterned prey harder to catch. Anim. Cogn. 18, 1059–1068. doi:10.1007/s10071-015-0874-0

Harland, D. P. and Jackson, R. R. (2000). Cues by which Portia fimbriata, an araneophagic jumping spider, distinguishes jumping-spider prey from other prey. J. Exp. Biol. 203, 3485–3494. doi:10.1242/jeb.203.22.3485

Harland, D. P., Jackson, R. R. and Macnab, A. M. (1999). Distances at which jumping spiders (Araneae: Salticidae) distinguish between prey and conspecific rivals. J. Zool. 247, 357–364. doi:10.1111/j.1469-7998.1999.tb00998.x

Harland, D. P., Li, D. and Jackson, R. R. (2012). How jumping spiders see the world. In How Animals See the World Comparative Behavior, Biology, and Evolution of Vision. (ed. O. F. Lazareva, T. Shimizu and E. Wasserman), pp. 133–164. Oxford, UK: Oxford University Press.

Hodos, W. (2012). What birds see and what they don’t. In How Animals See the World: Comparative Behavior, Biology, and Evolution of Vision (ed. O. F. Lazareva, T. Shimizu and E. Wasserman), pp. 5–24. Oxford, UK: Oxford University Press.

Ioannou, C. C. and Krause, J. (2009). Interactions between background matching and motion during visual detection can explain why cryptic animals keep still. Biol. Lett. 5, 191–193. doi:10.1098/rsbl.2008.0758

Jackson, R. and Blest, A. (1982). The biology of Portia fimbriata, a web-building jumping spider (Araneae, Salticidae) from Queensland: Utilization of webs and predatory versatility. J. Zool. 196, 255–293. doi:10.1111/j.1469-7998.1982.tb03504.x

Jackson, R. R. and Hallas, S. E. A. (1986). Comparative biology of Portia africana, P. Albimana, P. fimbriata, P. labiata, and P. shultzi, araneophagic, web-building jumping spiders (Araneae: Salticidae): Utilisation of webs, predatory versatility, and intraspecific interactions. N. Z. J. Zool. 13, 423–489. doi:10.1080/03014223.1986.10422978

Jakob, E. M., Long, S. M., Harland, D. P., Jackson, R. R., Carey, A., Searles, M. E., Porter, A. H., Canavesi, C. and Rolland, J. P. (2018). Lateral eyes direct principal eyes as jumping spiders track objects. Curr. Biol. 28, R1092–R1093. doi:10.1016/j.cub.2018.07.065

Jones, M. P., Pierce Jr, K. E. and Ward, D. (2007). Avian vision: a review of form and function with special consideration to birds of prey. J. Exot. Pet. Med. 16, 69–87. doi:10.1053/j.jepm.2007.03.012

Juanes, F. (1992). Why do decapod crustaceans prefer small-sized molluscan prey? Mar. Ecol. Prog. Ser. 87, 239–239. http://www.jstor.org/stable/24831652

Koh, J. K. H., Court, D. J., Ang, C. S. P. and Ng, P. Y. C. (2022). A Photographic Guide to Singapore Spiders. Singapore: National Parks Board.

Land, M. (1985). The morphology and optics of spider eyes. In Neurobiology of Arachnids (ed. F. G. Barth), pp. 53–78. Heidelberg, Berlin: Springer.

Li, D., Jackson, R. R. and Barrion, A. (1997). Prey preferences of Portia labiata, P. africana, and P. schultzi, araneophagic jumping spiders (Araneae: Salticidae) from the Philippines, Sri Lanka, Kenya, and Uganda. N. Z. J. Zool. 24, 333–349. doi:10.1080/03014223.1997.9518129

Maddison, W. P. (2015). A phylogenetic classification of jumping spiders (Araneae: Salticidae). J. Arachnol. 231–292. http://www.jstor.org/stable/24717307

Maddison, W. P. and Hedin, M. C. (2003). Jumping spider phylogeny (Araneae: Salticidae). Invertebr. Syst. 17, 529–549. doi:10.1071/IS02044

Michalis, C., Scott-Samuel, N. E., Gibson, D. P. and Cuthill, I. C. (2017). Optimal background matching camouflage. Proc. R. Soc. B. 284, 20170709. doi:10.1098/rspb.2017.0709

Mizutani, A., Chahl, J. S. and Srinivasan, M. V. (2003). Motion camouflage in dragonflies. Nature. 423, 604–604. doi:10.1038/423604a

Murali, G. and Kodandaramaiah, U. (2018). Body size and evolution of motion dazzle coloration in lizards. Behav. Ecol. 29, 79–86. doi:10.1093/beheco/arx128

Nakazawa, T., Ohba, S. Y. and Ushio, M. (2013). Predator–prey body size relationships when predators can consume prey larger than themselves. Biol. Lett. 9, 20121193. doi:10.1098/rsbl.2012.1193

Regan, D. and Beverley, K. (1984). Figure–ground segregation by motion contrast and by luminance contrast. JOSA A. 1, 433–442. doi:10.1364/JOSAA.1.000433

Rushton, S. K., Bradshaw, M. F. and Warren, P. A. (2007). The pop out of scene-relative object movement against retinal motion due to self-movement. Cognition. 105, 237–245. doi:10.1016/j.cognition.2006.09.004

Ruxton, G. D., Allen, W. L., Sherratt, T. N. and Speed, M. P. (2019). Avoiding attack: the evolutionary ecology of crypsis, aposematism, and mimicry. Oxford, UK: Oxford University Press.

Schneider, C. A., Rasband, W. S. and Eliceiri, K. W. (2012). NIH Image to ImageJ: 25 years of image analysis. Nat. Methods. 9, 671–675. doi:10.1038/nmeth.2089

Schmitz, O. (2017). Predator and prey functional traits: understanding the adaptive machinery driving predator–prey interactions. F1000Research. 6. doi:10.12688/f1000research.11813.1

Scott-Samuel, N. E., Baddeley, R., Palmer, C. E. and Cuthill, I. C. (2011). Dazzle camouflage affects speed perception. PLoS One. 6, e20233. doi:10.1371/journal.pone.0020233

Smart, I. E., Cuthill, I. C. and Scott-Samuel, N. E. (2020). In the corner of the eye: camouflaging motion in the peripheral visual field. Proc. R. Soc. B. 287, 20192537. doi:10.1098/rspb.2019.2537

Steinhoff, P. O., Uhl, G., Harzsch, S. and Sombke, A. (2020). Visual pathways in the brain of the jumping spider Marpissa muscosa. J. Comp. Neurol. 528, 1883–1902. doi:10.1002/cne.24861

Stevens, M. and Merilaita, S. (2011). Animal Camouflage: Mechanisms and Function. Cambridge, UK: Cambridge University Press.

Stevens, M., Párraga, C. A., Cuthill, I. C., Partridge, J. C. and Troscianko, T. S. (2007). Using digital photography to study animal coloration. Biol. J. Linn. Soc. 90, 211–237. doi:10.1111/j.1095-8312.2007.00725.x

Stevens, M., Yule, D. H. and Ruxton, G. D. (2008). Dazzle coloration and prey movement. Proc. R. Soc. B. 275, 2639–2643. doi:10.1098/rspb.2008.0877

Su, K., Meier, R., Jackson, R., Harland, D. and Li, D. (2007). Convergent evolution of eye ultrastructure and divergent evolution of vision-mediated predatory behaviour in jumping spiders. J. Evol. Biol. 20, 1478–1489. doi:10.1111/j.1420-9101.2007.01335.x

Troscianko, T., Benton, C. P., Lovell, P. G., Tolhurst, D. J. and Pizlo, Z. (2009). Camouflage and visual perception. Philos. Trans. R. Soc. Lond. B Biol. Sci. 364, 449–461. doi:10.1098/rstb.2008.0218

Umeton, D., Read, J. C. and Rowe, C. (2017). Unravelling the illusion of flicker fusion. Biol. Lett. 13, 20160831. doi:10.1098/rsbl.2016.0831

Umeton, D., Tarawneh, G., Fezza, E., Read, J. C. A. and Rowe, C. (2019). Pattern and Speed Interact to Hide Moving Prey. Curr. Biol. 29, 3109–3113 e3. doi:10.1016/j.cub.2019.07.072

Watanabe, H. and Yano, E. (2013). Behavioral response of mantid Tenodera aridifolia (Mantodea: Mantidae) to windy conditions as a cryptic approach strategy for approaching prey. Entomol. Sci. 16, 40–46. doi:10.1111/j.1479-8298.2012.00536.x

Yin, J., Gong, H., An, X., Chen, Z., Lu, Y., Andolina, I. M., McLoughlin, N. and Wang, W. (2015). Breaking cover: neural responses to slow and fast camouflage-breaking motion. Proc. R. Soc. B. 282, 20151182. doi:10.1098/rspb.2015.1182

Zurek, D. B., Taylor, A. J., Evans, C. S. and Nelson, X. J. (2010). The role of the anterior lateral eyes in the vision-based behaviour of jumping spiders. J. Exp. Biol. 213, 2372–2378. doi:10.1242/jeb.042382

